# DNA Break-Induced Epigenetic Alterations Promote Plaque Formation and Behavioral Deficits in an Alzheimer’s Disease Mouse Model

**DOI:** 10.1101/2025.10.21.683739

**Authors:** Sydney Bartman, Dakota Hunter, Lauren Gaspar, Hannah Tobias-Wallingford, Danielo Zamor, David A. Sinclair, Giuseppe Coppotelli, Jaime M. Ross

## Abstract

The dramatic increase in human longevity over recent decades has contributed to a rising prevalence of age-related diseases, including neurodegenerative disorders such as Alzheimer’s disease (AD). While accumulating evidence implicates DNA damage and epigenetic alterations in the pathogenesis of AD, their precise mechanistic role remains unclear. To address this, we developed a novel mouse model, DICE (Dementia from Inducible Changes to the Epigenome), by crossing the APP/PSEN1 (APP/PS1) transgenic AD model with the ICE (Inducible Changes to the Epigenome) model, which allows for the controlled induction of double-strand DNA breaks (DSBs) to stimulate aging-related epigenetic drift. We hypothesized that DNA damage induced epigenetic alterations could influence the onset and progression of AD pathology. After experiencing DNA damage for four weeks, DICE mice, together with control, ICE, and APP/PS1 mice, were allowed to recover for six weeks before undergoing a battery of behavioral assessments including the open-field test, light/dark preference test, elevated plus maze, Y-maze, Barnes maze, social interaction, acoustic startle, and pre-pulse inhibition (PPI). Molecular and histological analyses were then performed to assess Aβ pathology and neuroinflammatory markers. Our findings reveal that DNA damage-induced epigenetic changes significantly affect cognitive behavior and alters Aβ plaque morphology and neuroinflammation as early as six months of age. These results provide the first direct evidence that DNA damage can modulate amyloid pathology in a genetically susceptible AD model. Future studies will be aimed at investigating DNA damage– induced epigenetic remodeling across additional models of AD and neurodegeneration to further elucidate its role in brain aging and disease progression.

## Introduction

As the average human lifespan has dramatically increased over the last century, the prevalence of age-related disorders has also risen, with neurodegenerative diseases at the forefront (Crimmins, 2015). In 2024, approximately 7 million individuals in the US were diagnosed with a neurodegenerative disorder, with an estimated 6.5 million of them living with Alzheimer’s disease (AD) alone (“2024 Alzheimer’s disease facts and figures,” 2024). Given that the prevalence of AD in the US is projected to reach 13.8 million by 2060 (“2024 Alzheimer’s disease facts and figures,” 2024), understanding the underlying biological mechanisms has become imperative.

AD is a progressive neurodegenerative disorder primarily characterized by the accumulation of extracellular amyloid-beta plaques, also known as “senile plaques”, and intracellular neurofibrillary tangles formed from hyperphosphorylated tau protein, and is accompanied by widespread neurodegeneration (Breijyeh and Karaman, 2020). According to the “Amyloid Cascade Hypothesis”, the formation of amyloid-beta plaques is considered the earliest toxic event in the pathogenesis of AD (Kepp et al., 2023; Wu et al., 2022). These plaques form from the abnormal proteolytic cleavage of the amyloid precursor protein (APP) by the beta- and gamma-secretases, leading to the release of insoluble amyloid-beta oligomers that ultimately disrupt neuronal signaling and result in neuronal death (Chen et al., 2017). In the human brain, AD pathology typically begins in regions such as hippocampus and entorhinal cortex (Kobro-Flatmoen et al., 2021; Sheppard and Coleman, 2020), resulting in impaired learning and memory (Berry and Harrison, 2023; Blackmer-Raynolds et al., 2025). As the pathology spreads to additional brain areas, deficits in complex attention, language and executive, visuospatial, and social functioning progressively emerge (Long and Holtzman, 2019). Notably, the dominant familial form of AD, known as early onset AD (EOAD), that arises from mutation in one or more of three genes: presenilin 1 (*PSEN1*), presenilin 2 (*PSEN2*), and *APP*, only represents about 5% of cases, whereas sporadic AD, also known as late onset AD (LOAD), constitutes approximately 95% of cases and is influenced by a combination of genetic predispositions, environmental exposures, and aging-related factors (Andrade-Guerrero et al., 2023).

Epigenetic alterations consisting in changes in DNA methylation patterns, histone modifications, and chromatin structure are a recognized hallmark of aging (Gong et al., 2015; Kriukov et al., 2024; A. T. Lu et al., 2023; Shah et al., 2013; Xiao et al., 2016). These modifications lead to altered gene expression and compromised genome stability, contributing to cellular dysfunction and the progression of age-related diseases and have been implicated in AD etiology (Sanchez-Mut and Gräff, 2015). In this regard, DNA methylation, an epigenetic alteration consisting in the addition of a methyl group to the guanosine base of DNA that has been linked to gene repression, has been found to become altered in AD. Specifically, one study identified 334 differentially methylated DNA sites (DMPs) in the cortex of the human brain associated with AD pathology, while another study analyzing DNA methylation in APP/PS1 transgenic mice revealed 485 hypermethylated genes (Cong et al., 2014; Shireby et al., 2022). In addition to DNA methylation, histone acetylation, an epigenetic mechanism in which an acetyl group is added to lysine residues on histone proteins, has also been found to be dysregulated in AD. For instance, Xu et al. (2024) reported that H3K27ac may contribute to mitigating AD pathology in CNS neurons by promoting the expression of compensatory genetic programs (Xu et al., 2024). Similarly, Nativio et al. (2018) identified a global loss of H4K16ac near genes associated with aging and AD in human brain tissue (Nativio et al., 2018).

Importantly, these epigenetic alterations are driven by a complex interplay of genetic and environmental factors, including age, lifestyle, and underlying genetic predispositions. As a result, the mechanisms underlying these changes and their contribution to disease progression and pathology are still under investigation. DNA damage, defined as any alteration to the chemical structure or integrity of the DNA molecule, including base modification, strand breaks, and crosslinks, has also been implicated in AD, although its mechanistic role remains poorly understood (Fernandez et al., 2021; Lin et al., 2020). DNA damage can disrupt key cellular processes such as DNA repair, transcription, DNA damage response (DDR) signaling, and epigenomic stability (Hakem, 2008; Manfrini et al., 2015; Mendes et al., 2024; Nikfarjam and Singh, 2023; Park et al., 1999). Evidence linking DNA damage to AD comes from both human postmortem studies and experimental models. For example, Zhang et al. (2023) reported elevated levels of *γ*-H2AX, a marker of double-strand DNA breaks, in postmortem brain tissue from individuals with AD (Zhang et al., n.d.), and Foret et al. (2024) similarly observed higher levels of *γ*-H2AX in brains from APP/PS1 mice (Foret et al., 2024). In addition, Jiang et al. (2021) found significantly higher levels of 8-hydroxy-2’-deoxyguanosine (8-OHdG), a marker of oxidative DNA damage, in APP/PS1 mice (Jiang et al., 2021). Despite these associations, it remains unclear whether amyloid-beta plaque formation drives, or is driven by, DNA damage and its associated epigenetic alterations.

To address this question, we investigated the impact of DNA damage–induced epigenetic alterations on amyloid-beta plaque pathology and neuroinflammation in a mouse model of AD. Specifically, we combined the well-characterized APP/PS1 transgenic mouse model (Jankowsky et al., 2004) with the “ICE” (Inducible Changes to the Epigenome) model developed by Oberdoerffer and Sinclair, which induces site-specific double-strand DNA breaks (DSBs) to mimic age-associated epigenetic drift (Kim et al., 2016; Yang et al., 2023). The model was designed to test the relocalization of chromatin modifiers (RCM) hypothesis, which posits that DNA double-strand breaks recruit chromatin modifiers (e.g., SIRT1, SIRT6, HDAC1) to sites of repair, causing epigenetic information loss when they fail to return to their original loci (Oberdoerffer et al., 2008).

The cross resulted in a novel model, termed “DICE” (Dementia from Inducible Changes to the Epigenome). The ICE system utilizes the I-*Ppo*I endonuclease derived from *Physarum polycephalum* to induce lower level of DSBs throughout the mouse genome (Kim et al., 2016). Upon tamoxifen administration, CRE-ER^T2^ recombinase, expressed ubiquitously under the control of a ubiquitin promoter (“007001 - UBC-Cre-ERT2 Strain Details,” n.d.; Feil et al., 1997, 1996), becomes activated and translocates into the nucleus, where it excises the STOP codon cassette located upstream of the HA-ER^T2^-I-*Ppo*I-IRES-GFP coding sequence (**Figure 1A**). As a result of this, the HA-ER^T2^-I-*Ppo*I fusion protein is expressed and in the presence of tamoxifen relocalizes into the nucleus and cleaves the DNA at the recognized sequence CTCTCTT▼AAGGTAGC. This sequence is found at 20 canonical sites in the murine genome, of which 19 are noncoding regions, and none are present in the mitochondrial DNA (Kim et al., 2016). Using the ICE model, we together with our collaborators demonstrated that cycles of site-specific DSBs and repair caused by a tamoxifen-induced I-*Ppo*I, accelerated physiological, cognitive, and molecular markers of aging in mice (Yang et al., 2023). This was accompanied by widespread erosion of the epigenetic landscape, including loss of cell identity (dedifferentiation), activation of senescence-associated gene signatures, and acceleration of the DNA methylation age clock (Yang et al., 2023). This study extends the RCM (Oberdoerffer and Sinclair, 2007) and the Information Theory of Aging (Y. R. Lu et al., 2023), which propose that DNA damage–induced relocalization of chromatin modifiers erodes epigenetic information and cell identity over time. By applying this framework to Alzheimer’s pathology, we tested whether induced epigenetic noise synergizes with amyloidogenic stress to accelerate disease, specifically the formation, spatial distribution, and neuroinflammatory response associated with amyloid-beta plaques.

**Figure 1.**
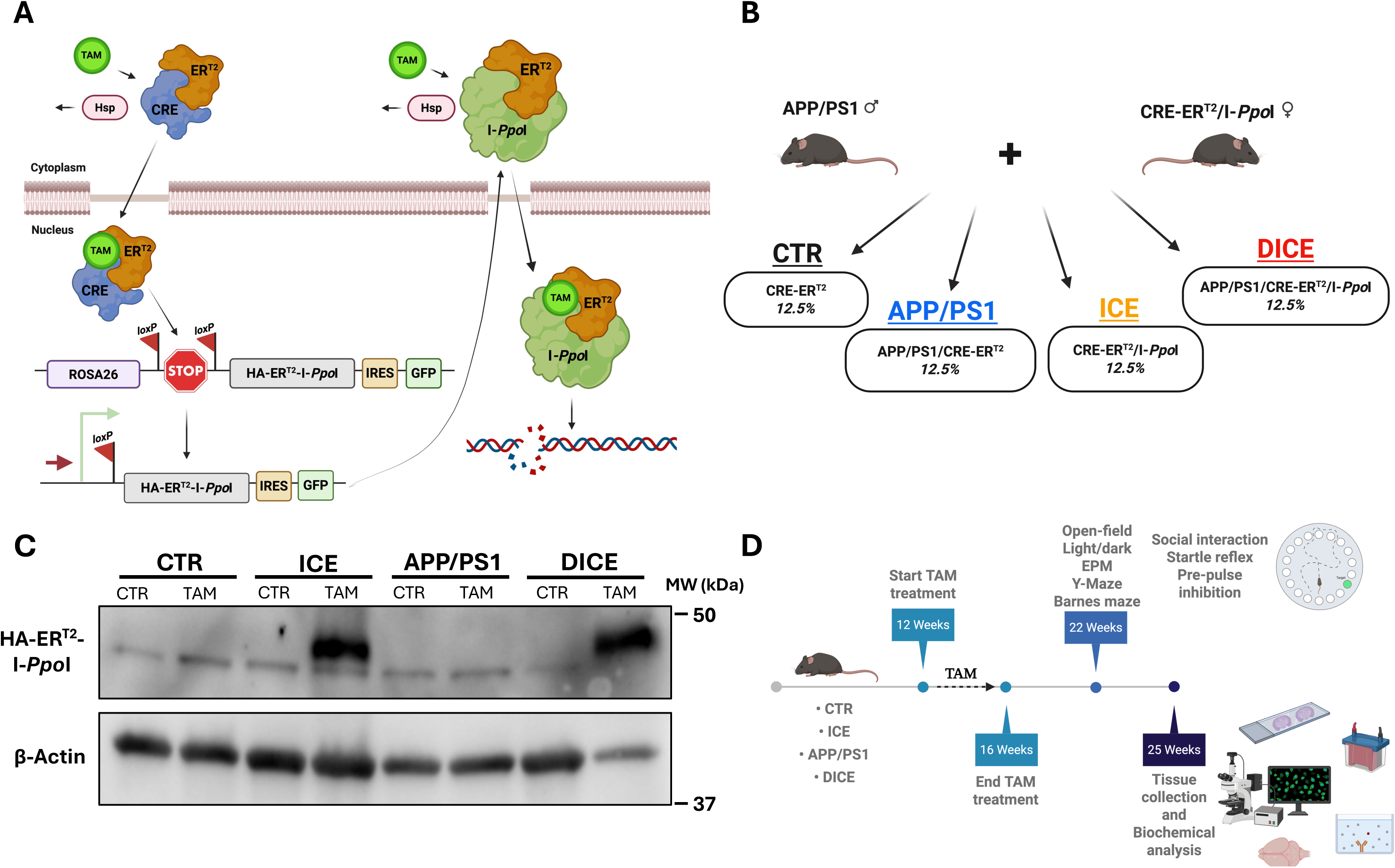
DICE mouse system and experimental timeline. **(A)** Schematic of the ICE system. Upon tamoxifen binding, the Cre-ER^T2^ protein excises the STOP cassette, enabling expression of HA-ER^T2^-I-*Ppo*I and GFP proteins. Tamoxifen allows HA-ER^T2^-I-*Ppo*I translocation into the nucleus, where it induces double-strand breaks (DSBs) throughout the genome without causing mutations or aneuploidy. **(B)** Breeding strategy used to generate experimental mouse groups. **(C)** Representative Western blot confirming HA-ER^T2^-I-*Ppo*I expression in ICE and DICE fibroblasts treated with 0.5 µM tamoxifen citrate. (**D)** Schematic of the experimental timeline. Panels A, B, C were created with BioRender (accessed on 31 August 2025).

## Results

Female and male hemizygous DICE mice were generated by crossing hemizygous HA-ER^T2^-I-*Ppo*I/Cre-ER^T2^ females with APP/PS1 males (**Figure 1B**). Since Cre expression alone did not affect mouse phenotype (data not shown), experimental groups included CTR (Cre-ER^T2^ or WT), APP/PS1 (APP/PS1 or APP/PS1/Cre-ER^T2^), ICE, and DICE genotypes to ensure the ethical use of all offspring. To confirm proper activation of the ICE system, primary fibroblasts were isolated from each genotype and treated *in vitro* with 0.5 µM 4-hydroxytamoxifen for 72 hours to assess HA-ER^T2^-I-*Ppo*I expression (**Figure 1C**). From 12 to 16 weeks of age, CTR, APP/PS1, ICE, and DICE mice were treated with tamoxifen (360 mg/kg in food) to induce DNA damage by promoting the nuclear translocation of ER^T2^-I-*Ppo*I following Cre-ER^T2^ mediated excision of the STOP cassette in the ICE and DICE models. At 22 weeks of age, mice underwent a battery of behavioral assays, after which they were sacrificed at 25 weeks (∼6 months) of age, and tissues were collected for histological and biochemical analyses (**Figure 1D**).

### I-PpoI Recognition Sites

To ensure that the HA-ER^T2^-I-*Ppo*I restriction enzyme did not recognize or cleave any site within the human APP coding sequence (CDS), HA-tagged I-*Ppo*I endonuclease was expressed in U-2 OS cells and isolated by immunoprecipitation using HA agarose beads (**Figure 2A**). Following isolation, HA-I-*Ppo*I was incubated either with the p201B/Cas9 plasmid containing an I-*Ppo*I recognition site as a positive control (Jacobs et al., 2015) or with purified hu-APP CDS amplified by PCR from olfactory bulbs from APP/PS1 mice. While I-*Ppo*I efficiently cleaved the Cas9 plasmid at the expected site, no cleavage was observed in the isolated hu-APP CDS, suggesting that I-*Ppo*I does not recognize any non-canonical site within this sequence *in vitro* (**Figure 2B**). To further verify that *in vivo* activated I-*Ppo*I does not cleave the hu-APP sequence at any site, PCR-amplified hu-APP products in olfactory bulbs from APP/PS1, and DICE with and without tamoxifen treatment were analyzed by Sanger sequencing, confirming the absence of any mutations potentially caused by DNA damage (**Figure S1**). To verify that activated I-*Ppo*I endonuclease does not affect hu-APP mRNA expression or protein levels, total mRNA and proteins were extracted from hemi-cerebellum and olfactory bulbs from CTR, APP/PS1, and DICE mice, and hu-APP mRNA and protein levels were measured by qPCR and Western blot analyses, respectively. As expected, no hu-APP mRNA or protein expression was detected in CTR mice, while no significant differences in hu-APP mRNA or protein levels were observed between DICE and APP/PS1 mice (**Figure 2C, D, E**).

**Figure 2.**
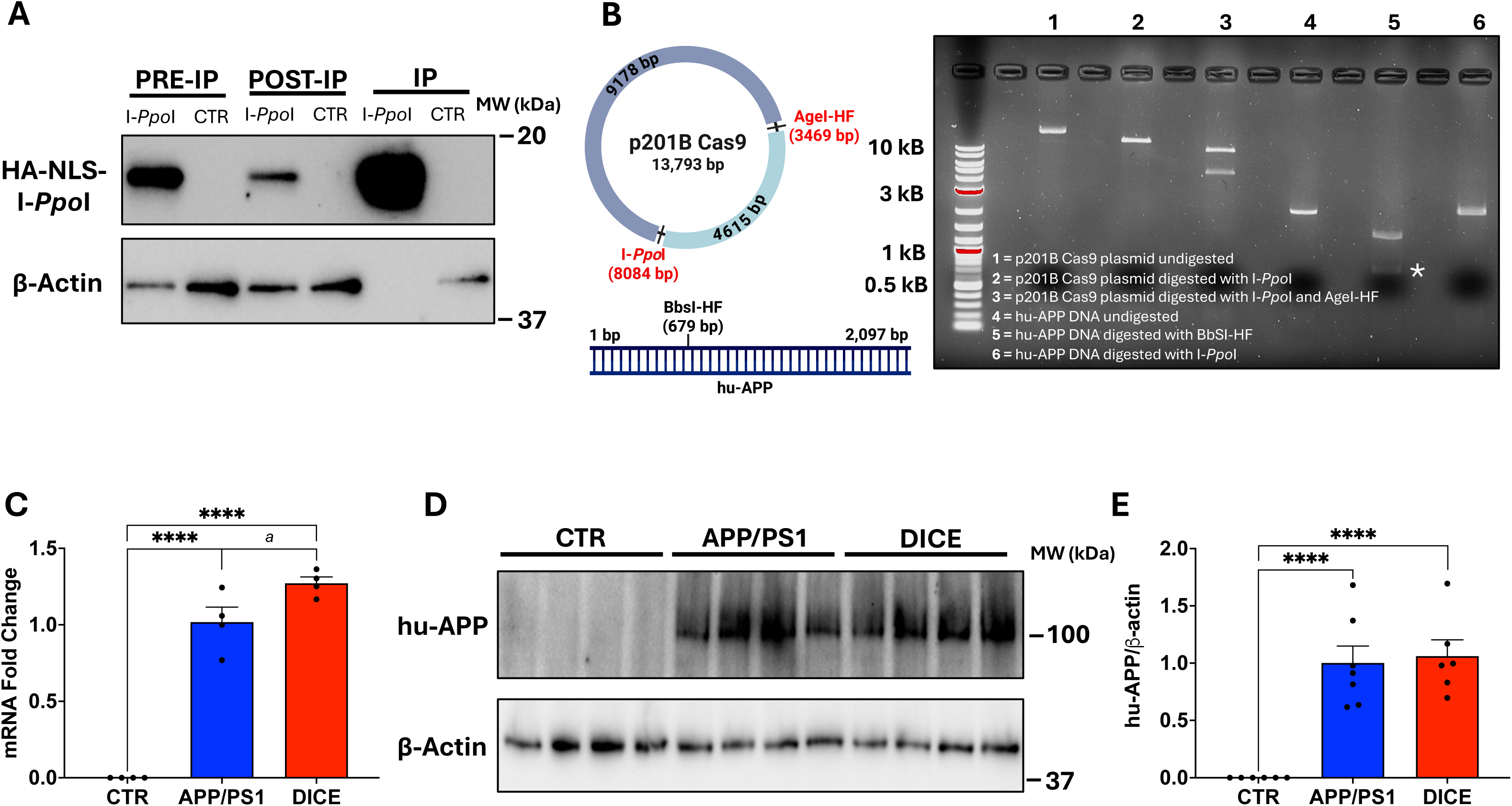
The humanized APP gene lacks recognition sites for I-*Ppo*I endonuclease. **(A)** Western blot analysis showing expression and immunoprecipitation of native HA-I-*Ppo*I from transfected U-2 OS cells. **(B)** Schematic representation of the p201B/Cas9 plasmid and hu-APP sequences, indicating the positions of I-*Ppo*I, AgeI, and BbsI restriction sites. An agarose gel electrophoresis image is shown for p201B/Cas9 (positive control) and hu-APP PCR products digested with the indicated enzymes. **(C)** Quantitative PCR analysis of hu-APP mRNA levels in APP/PS1 and DICE mice compared to control (CTR) mice (N=4 per group). **(D, E)** Representative Western blot and corresponding quantification of hu-APP protein in brain tissue from CTR, APP/PS1, and DICE mice (N=6-7 per group). Data are presented as mean ± SEM and analyses were performed using unpaired t-test with Welch’s correction: *a* indicates a trend p<0.10 and ****p<0.0001.

### Phenotype of DICE Mice

Since ICE mice were previously reported to exhibit a premature aging phenotype 10 months after tamoxifen treatment (Yang et al., 2023), all experimental groups were evaluated before behavioral testing for signs of premature aging using the frailty index scoring system (Whitehead et al., 2014).

Six weeks after tamoxifen treatment, male and female ICE and DICE mice start to exhibit signs of premature aging, including hair greying, cataract formation, and loss of skin pigmentation (**Figure 3A**). While body weight was reduced in both ICE and DICE mice compared to CTR, regardless of sex, only female DICE mice weighed less compared to APP/PS1 mice (**Figure 3B**). When mice were assessed for frailty index (FI), both female ICE and DICE mice had markedly higher FI scores compared to CTR and APP/PS1 mice, whereas in males, this difference was only in the ICE group (**Figure 3C**). Finally, we observed no differences in normalized brain weight among groups (**Figure 3D**).

**Figure 3.**
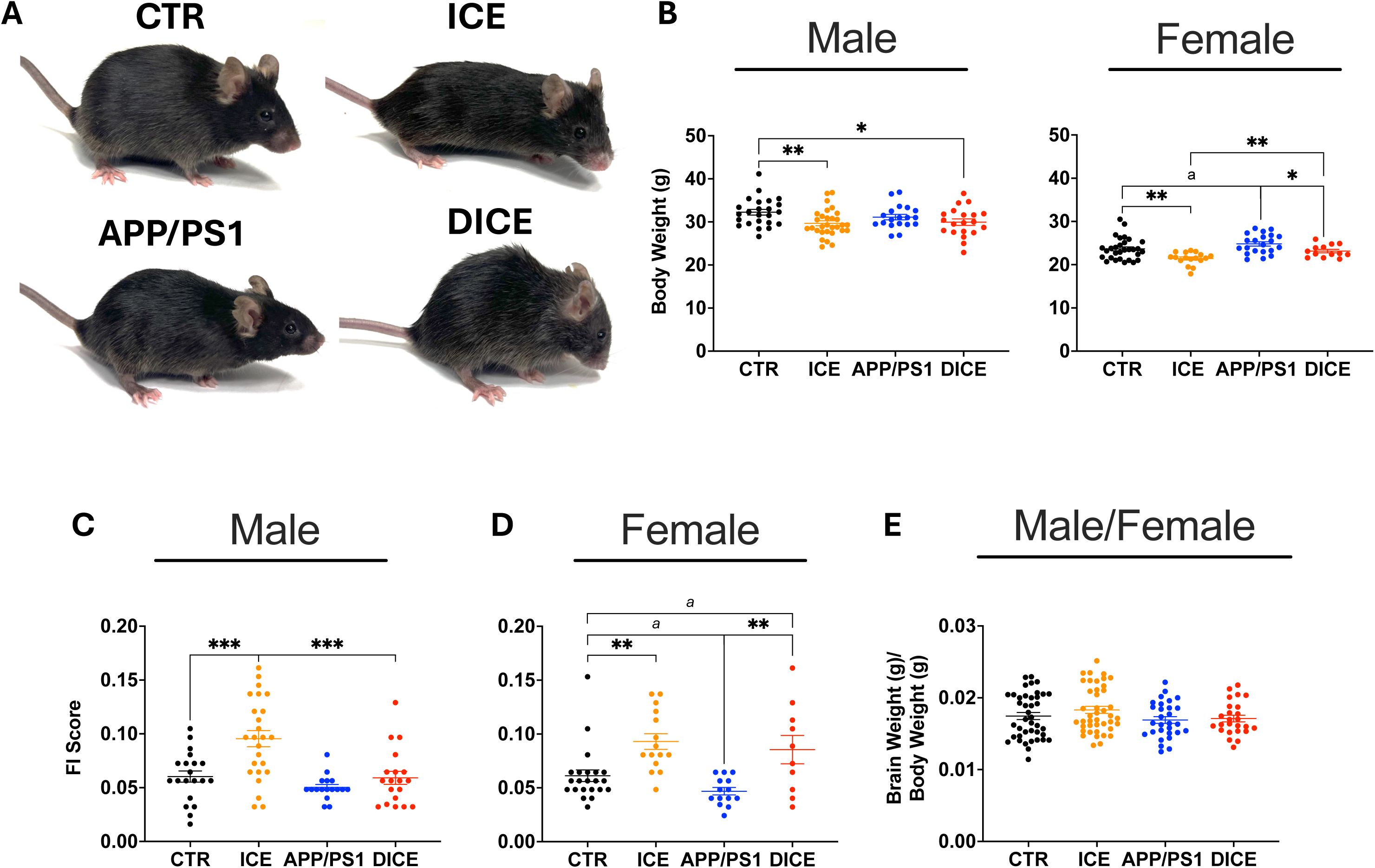
DICE mice exhibit signs of premature aging compared to control groups. **(A)** Representative images of CTR, ICE, APP/PS1, and DICE mice at 25 weeks of age. **(B)** Body weight measurements of male and female mice at 25 weeks of age weights weights [CTR (N_Male_=24; N_Female_=30), ICE (N_Male_=30; N_Female_=17), APP/PS1 (N_Male_=19; N_Female_=21), and DICE (N_Male_=20; N_Female_=13)]. (**C, D)** Frailty Index (FI) scores assessed at 25 weeks of age index [CTR (N_Male_=20; N_Female_=22), ICE (N_Male_=25; N_Female_=15), APP/PS1 (N_Male_=17; N_Female_=14), and DICE (N_Male_=19; N_Female_=10)]. **(E)** Brain weight normalized to body weight [CTR (N_Male_=21; N_Female_=20), ICE (N_Male_=28; N_Female_=14), APP/PS1 (N_Male_=16; N_Female_=13), and DICE (N_Male_=18; N_Female_=7)]. Data are presented as mean ± SEM. Statistical analyses were performed using an unpaired t-test with Welch’s correction: *a* indicates a trend p<0.10, \**p*<0.05, \*\****p***<0.01, and *\*\*\***p***<0.001.

### Changes in Behavioral Correlates in DICE Mice

Six weeks post-tamoxifen treatment, behavioral analyses were conducted in order of increasing invasiveness: open-field, light/dark preference, elevated plus maze (EPM), Y-Maze, Barnes maze, social interaction, startle reflex, and pre-pulse inhibition. In the open-field test, both male and female DICE mice traveled significantly farther than APP/PS1, ICE, and CTR mice. APP/PS1 mice also traveled farther than ICE and CTR mice, thus supporting that the DNA damage exacerbated the increased hyperactivity observed in DICE mice (**Figure 4A**). Similarly, male and female DICE mice reared more as compared to age-matched APP/PS1, ICE, and CTR mice (**Figure 4B**). Both DICE and APP/PS1 mice spent markedly less time resting throughout the test, as well as in total, compared to CTR and ICE mice (**Figure 4C**). Likewise, male and female DICE and APP/PS1 mice reached higher peak average velocities during the assay compared to CTR and ICE mice, with DICE mice displaying significantly faster average velocities than CTR and ICE mice (**Figure 4D**).

**Figure 4.**
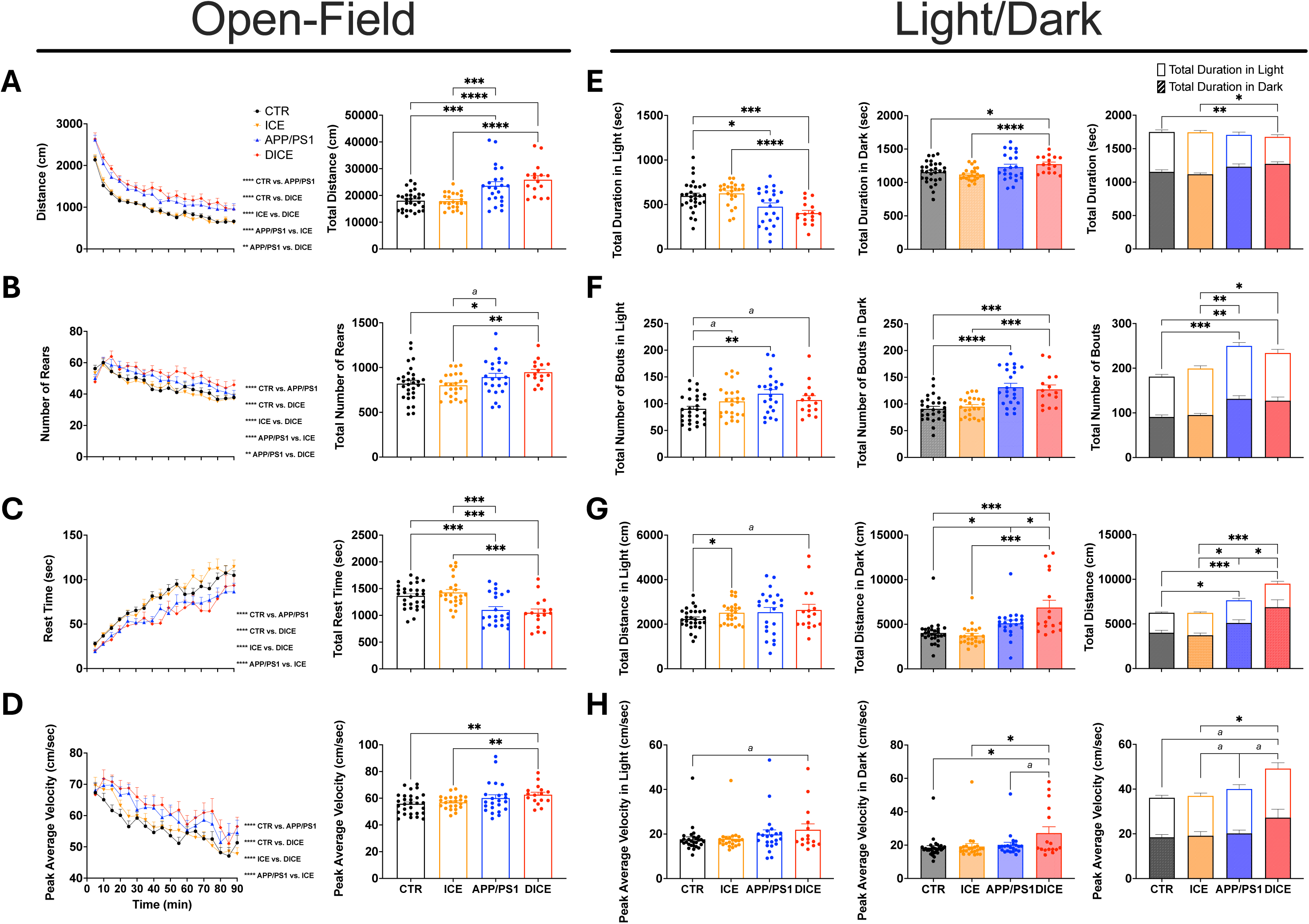
Open-field and light/dark transition tests reveal altered locomotor and anxiety-like behaviors in DICE mice following tamoxifen treatment. **(A)** Distance traveled, **(B)** number of rears, **(C)** rest time, and **(D)** peak average velocity of CTR, ICE, APP/PS1, and DICE mice assessed in the open-field test [CTR (N_Male_=13; N_Female_=16), ICE (N_Male_=14; N_Female_=10), APP/PS1 (N_Male_=13; N_Female_=10), and DICE (N_Male_=11; N_female_=5)]. **(E)** Total duration spent in the light and dark compartments [CTR (N_Male_=13; N_Female_=16), ICE (N_Male_=14; N_Female_=10), APP/PS1 (N_Male_=13; N_Female_=10), and DICE (N_Male_=11; N_Female_=5)], **(F)** total number of bouts in light and dark compartments [CTR (N_Male_=12; N_Female_=16), ICE (N_Male_=13; N_Female_=10), APP/PS1 (N_Male_=12; N_Female_=10), and DICE (N_Male_=10; N_Female_=5)] and **(G)** total distance traveled and peak average velocity in the light and dark compartments, assessed by the light/dark transition test [CTR (N_Male_ =12; N_Female_=16), ICE (N_Male_=13; N_Female_=10), APP/PS1 (N_Male_=12; N_Female_=10), and DICE (N_Male_=11; N_Female_=5)]. Data are presented as mean ± SEM. Statistical analyses were performed using two-way ANOVA with Tukey’s post hoc comparison or unpaired t-test with Welch’s correction: *a* indicates a trend p<0.10, \**p*<0.05, \*\**p*<0.01, \*\*\**p*< 0.001, and \*\*\*\**p*< 0.0001.

In the light/dark preference test, both DICE and APP/PS1 mice showed increased anxiety-like behavior, spending less time in the light and more time in the dark compared to ICE and CTR mice, with the effect being significant for DICE mice (**Figure 4E**). Both APP/PS1 and DICE mice exhibited more bouts in the dark chamber and traveled farther and faster overall, with these behaviors being more pronounced in DICE mice compared to APP/PS1 (**Figures 4F–H**). EPM analysis further confirmed increased anxiety-like behavior in both APP/PS1 and DICE mice compared to CTR and ICE mice, as evidenced by the reduced percentage of time spent and distance traveled in the open arms (**Figure 5A–B**). No differences in sociability were observed among the groups in the social interaction test (**Figure 5C**). Similarly, assessment of spatial learning and working memory using the Y-Maze assay revealed no significant differences in the number of entries into the novel arm among any of the groups tested (**Figure S2A**). In contrast, assessment of spatial learning and memory using Barnes Maze revealed that six-month-old male and female DICE and APP/PS1 mice spent significantly less time at the target location compared to their CTR counterparts, indicating impairments in spatial memory. Additionally, APP/PS1 mice spent significantly less time at the target location than ICE mice (**Figure 5D, Figure S2B**). Interestingly, DICE mice exhibited a much greater startle response to an acoustic stimulus compared to APP/PS1 and ICE mice. Both DICE and APP/PS1 mice also showed significantly higher startle responses than CTR mice, further suggesting an elevated excitability (**Figure 5E**). Finally, sensorimotor gating was assessed using the pre-pulse inhibition (PPI) test and both male and female DICE mice exhibited a significantly stronger baseline startle response compared to APP/PS1 mice, suggesting impairments in their sensory gating system (**Figure 5F**).

**Figure 5.**
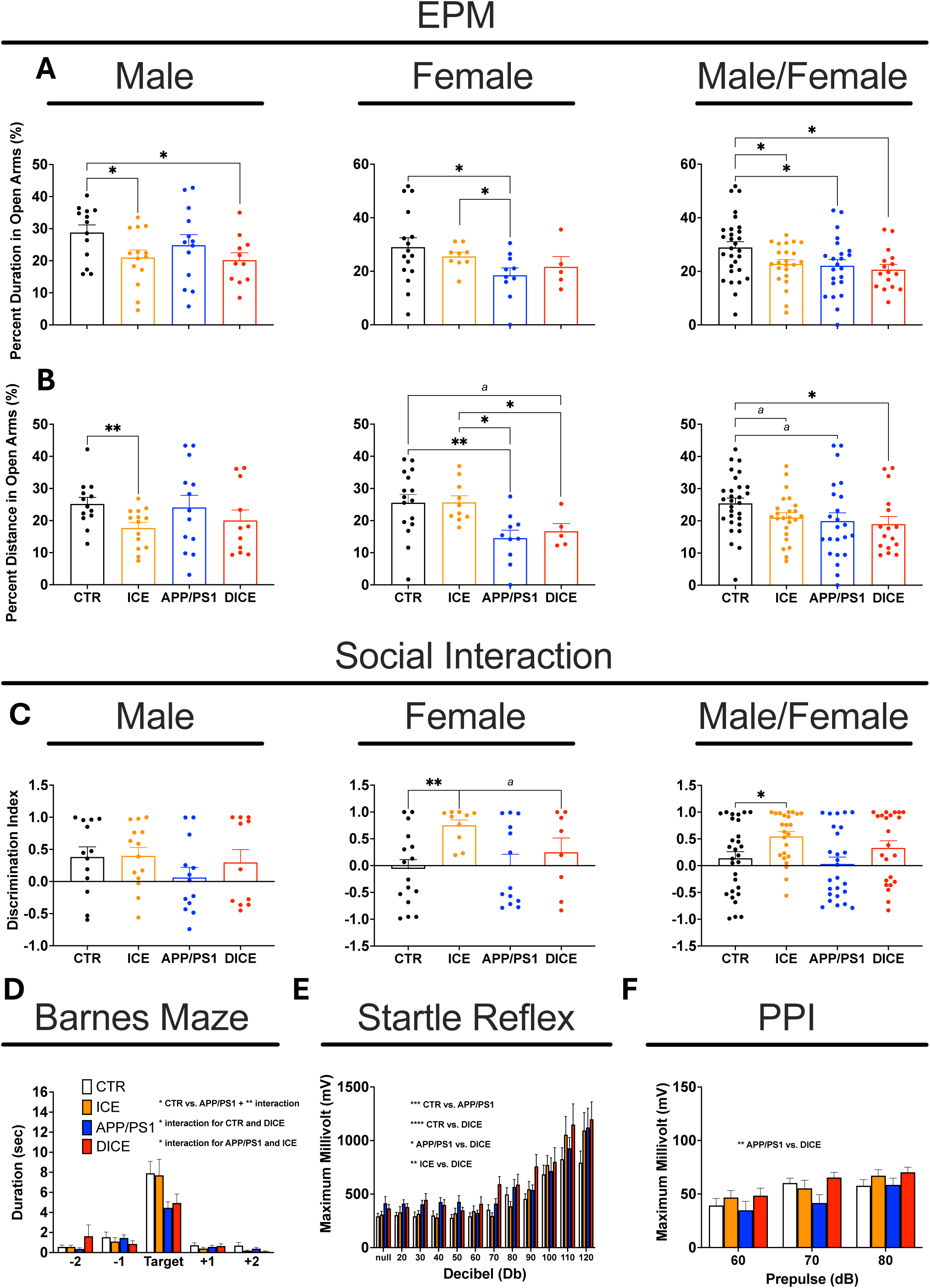
DICE mice exhibit heightened anxiety and startle responses with intact social and spatial behaviors. **(A)** Percent duration and **(B)** percent distance traveled in the open arms of the elevated plus maze [CTR (N_Male_=13; N_Female_=16), ICE (N_Male_=14; N_Female_=10), APP/PS1 (N_Male_=13; N_Female_=10), and DICE (N_Male_=11; N_Female_=5)]. **(C)** Social interaction ratio across genotypes [CTR (N_Male_=13; N_Female_=16), APP/PS1 (N_Male_=13; N_Female_=10), ICE (N_Male_=14; N_Female_=10), and DICE (N_Male_=10; N_Female_=5)]. **(D)** Duration in the target hole of the Barnes maze during the probe trial [CTR (N_Male_=9; N_Female_=15), ICE (N_Male_=10; N_Female_=10), APP/PS1 (N_Male_=10; N_Female_=8), and DICE (N_Male_=11; N_Female_=5)]. **(E)** Startle response to acoustic stimulus in the startle reflex paradigm [CTR (N_Male_=13; N_Female_=16), ICE (N_Male_=14; N_Female_=10), APP/PS1 (N_Male_=13; N_Female_=10),and DICE (N_Male_=11; N_Female_=5)]. **(F)** Pre-pulse inhibition response [CTR (N_Male_=11; N_Female_=11), ICE mice (N_Male_=16; N_Female_=6), APP/PS1 (N_Male_=8; N_Female_=6), DICE (N_Male_=5; N_Female_=5)]. Data represent male and female CTR, ICE, APP/PS1, and DICE mice post-tamoxifen treatment. Values are presented as mean ± SEM. Statistical analyses were performed using two-way ANOVA with Tukey’s post hoc comparison or unpaired t-test with Welch’s correction: *a* indicates a trend p<0.10, \**p*<0.05, \*\**p*<0.01, \*\*\**p*<0.001, \*\*\*\**p*<0.0001.

### Alterations in Histological Analyses in DICE Mice

To assess the effect of DNA damage–induced epigenetic drift on plaque load, amyloid-beta plaques were detected and quantified in CTR, ICE, APP/PS1 and DICE mice at 25 weeks of age using fluorescent immunohistochemistry and Thioflavin S staining. Six-month-old male and female hemi-sected brains from each group were cryosectioned and analyzed. As expected, DICE and APP/PS1 brain contained amyloid-beta plaques in areas such as cortex and hippocampus while CTR and ICE brains did not have any plaques (**Figure 6A and Figure S4A).** Quantification of plaque numbers in hippocampal sections showed that DICE mice had a higher median load compared to APP/PS1 mice; however, this difference was not statistically significant **(Figure 6B).** To further investigate plaque morphology in DICE and APP/PS1 mice, plaque area and perimeter were quantified. Interestingly, this analysis revealed significant differences between the groups with DICE mice having a higher number of plaques with smaller area and perimeter compared to APP/PS1 mice (**Figure 6C, D**). To confirm that tamoxifen administration did not affect amyloid-beta plaque formation or localization, hemi-sected brains from six-month-old male and female CTR, ICE, APP/PS1, and DICE mice that had not received tamoxifen were also analyzed (**Figure S3**).

**Figure 6.**
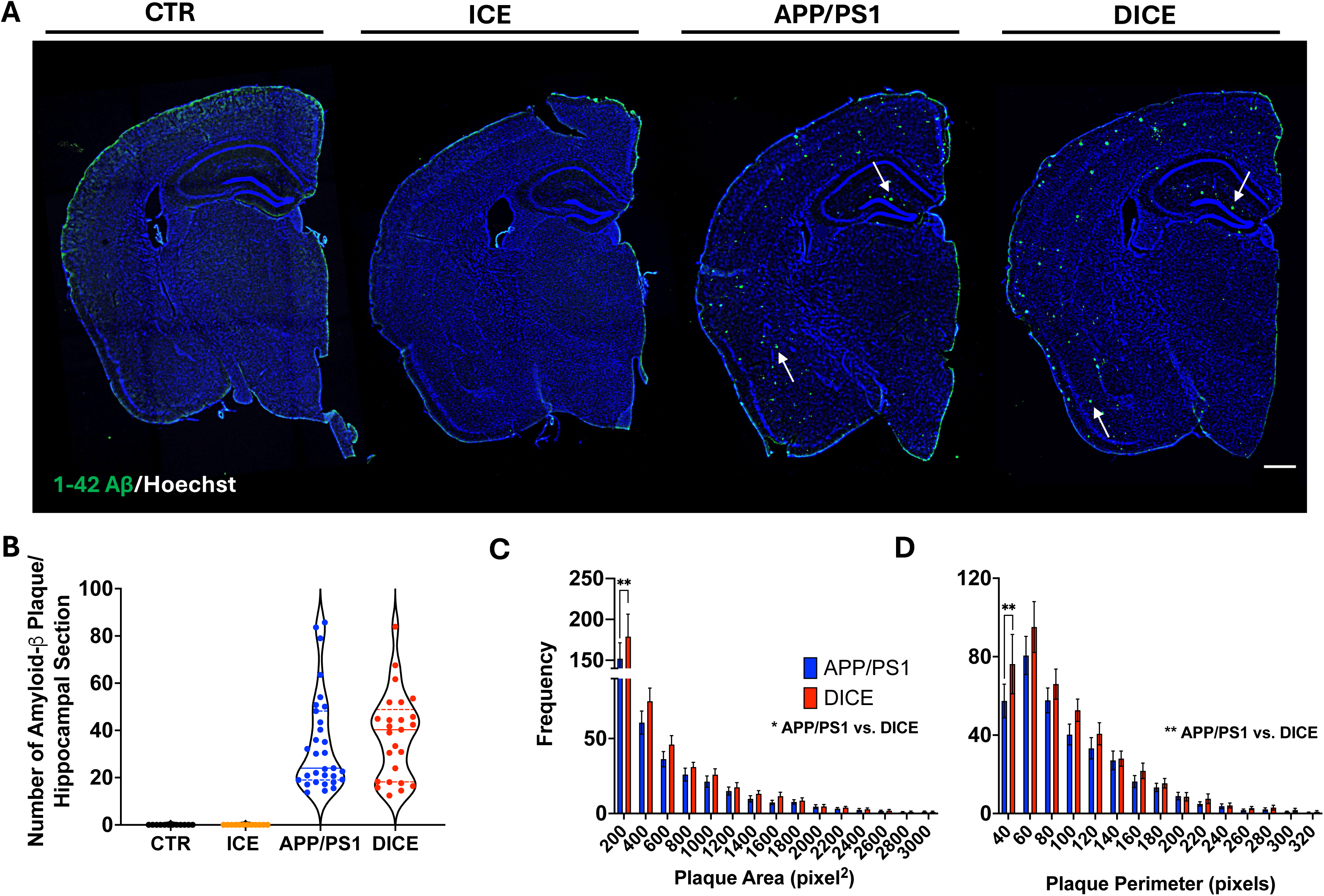
DICE mice exhibit altered amyloid-beta plaque morphology despite similar plaque load to APP/PS1 mice. **(A)** Representative fluorescent immunohistochemistry for Aβ1-42 and Hoechst in coronal brain sections from 6-month-old CTR (N=12), ICE (N=12), APP/PS1 (N=31), and DICE (N=25) mice. Significant plaque formation is observed in the hippocampus and cortex of APP/PS1 and DICE mice, with no plaques detected in CTR and ICE mice (5X; scale bar=500 μm). Arrows indicate examples of amyloid-beta plaques. **(B)** Manual quantification of Aβ1-42 plaques shows higher median plaque load in DICE mice as compared to APP/PS1, but no significant difference in average amounts was found. Frequency distributions of plaque area **(C)** and perimeter **(D)** reveal a higher proportion of smaller plaques in DICE brains compared to APP/PS1. Data are presented as mean ± SEM. Statistical analyses were performed using unpaired t-test with Welch’s correction: \**p* < 0.05 and \*\**p* < 0.01.

Following amyloid-beta plaque quantification, we examined overall brain morphology and assessed inflammation by immunostaining for glial fibrillary acidic protein (GFAP), a marker of activated astrocytes, and ionized calcium-binding adaptor molecule 1 (IBA1), a marker of activated microglia. Using cresyl violet staining, we found no gross changes in morphology or evidence of pyknotic bodies in any of the groups (**Figure S4B**). Interestingly, both DICE and APP/PS1 mice exhibited significantly higher diffuse GFAP and IBA1 expression in hippocampal and cortical regions, as compared to ICE and CTR mice (**Figure 7A**), with strong localization around plaques in these regions (data not shown). To better quantify changes in GFAP expression, protein levels were analyzed by Western blot in hemi-cerebellar lysates from CTR, ICE, APP/PS1, and DICE mice. This analysis confirmed elevated GFAP levels in DICE and APP/PS1 mice compared to CTR mice. Notably, DICE mice exhibited markedly higher GFAP protein level than APP/PS1 mice, indicating that DNA damage exacerbates astrocytic activation and inflammation in the presence of amyloid plaques (**Figure 7B–C**).

**Figure 7.**
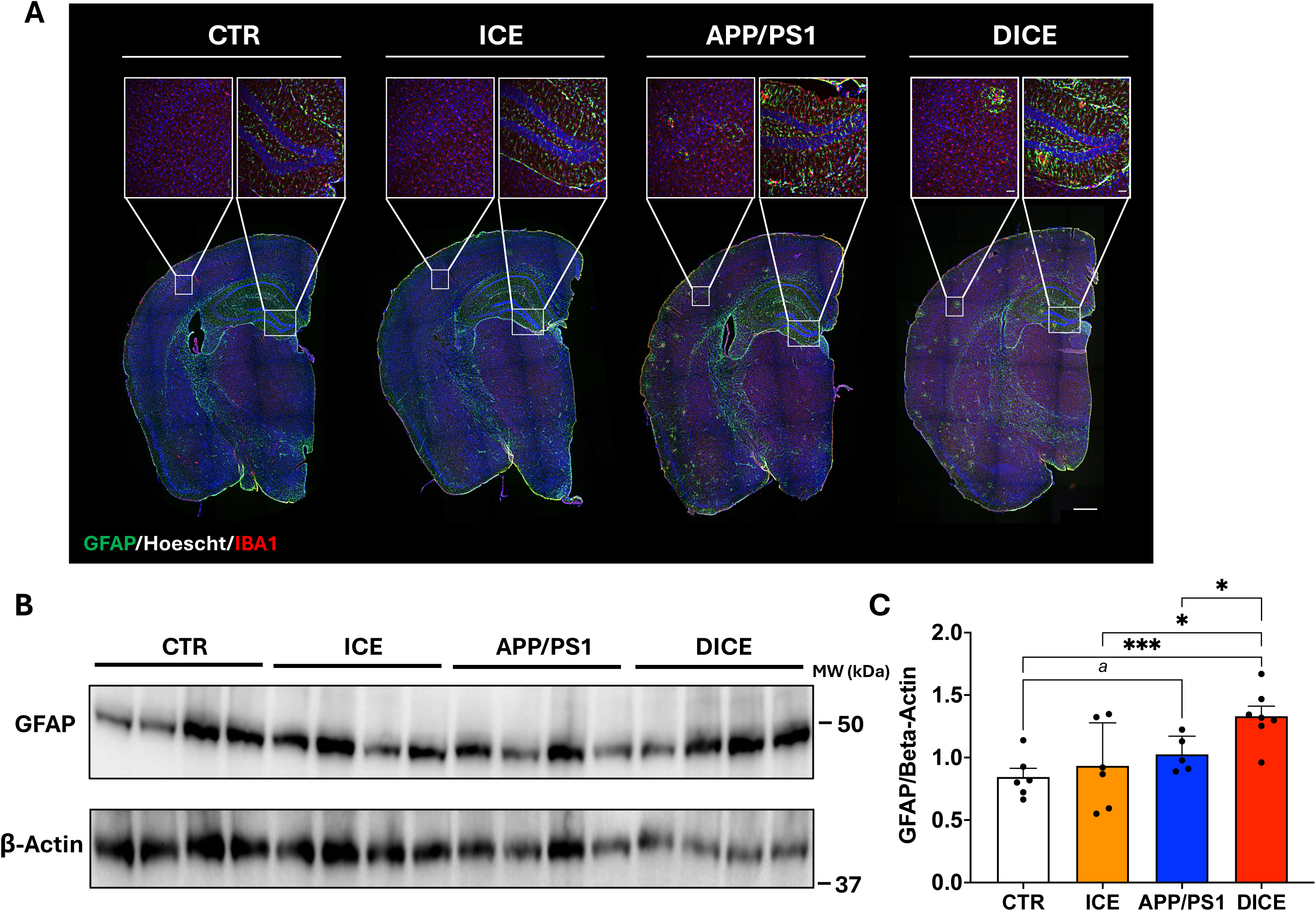
DICE mice exhibit increased gliosis compared to APP/PS1 and control mice. **(A)** Representative fluorescent immunohistochemistry for GFAP, IBA1, and Hoechst in coronal brain sections with cortex and dentate gyrus R.O.I.s, from 6-month-old CTR, ICE, APP/PS1, and DICE mice (N=3 per group). DICE and APP/PS1 mice show elevated astrocyte and microglial activation relative to ICE and CTR mice (brain: 5X, scale bar=500 μm; cortex and dentate gyrus: 20X, scale bar=50 μm). **(B)** Representative immunoblot and **(C)** quantification of cerebellar lysates show increased GFAP in 6-month-old DICE mice (N=7) compared to APP/PS1 (N=5), ICE (N=6), and CTR (N=6) mice. Data are presented as mean ± SEM and analyses were performed using unpaired t-test with Welch’s correction: *a* indicates a trend, p<0.10 *p<0.05, and ***p<0.001.

## Discussion

Aging is the strongest risk factor for Alzheimer’s disease (AD), and accumulating evidence suggests that loss of epigenomic integrity is a key mediator linking age to neurodegeneration (Lardenoije et al., 2015; Lu et al., 2004; Madabhushi et al., 2014). Using the DICE model, an APP/PS1 background in which low level, site-specific double-strand DNA breaks (DSBs) induce an aging-like epigenetic drift (Yang et al., 2023), we provide evidence that DNA damage alters amyloid pathology, neuroinflammation, and behavior early in the disease. These findings complement and extend prior reports that DNA damage and epigenetic dysregulation are pervasive in AD brains and models, but causality and directionality have remained difficult to disentangle (Lin et al., 2020).

Behaviorally, DICE mice exhibited exacerbated hyperactivity and anxiety-like phenotypes relative to APP/PS1 mice, as well as increased acoustic startle responses, whereas sociability and Y-maze performance remained largely unchanged. Findings of hyperactivity and anxiety-like behavior phenotypes in APP/PS1 mice have been controversial with some studies reporting increase in the APP/PS1 model while other found no change across the life span of the mouse (Huang et al., 2016; Locci et al., 2021; Webster et al., 2013). Here, we report that in our conditions, APP/PS1 mice exhibit hyperactivity and anxiety-like behavior early in the disease state, which is exacerbated by DNA damage in the DICE model. This aligns with recent work showing that persistent neuronal DSBs perturb gene programs regulating synaptic plasticity and excitability, potentially lowering thresholds for behavioral alterations (Konopka and Atkin, 2022).

A key finding of this study is the identification of altered plaque morphology, rather than overall plaque burden, in DICE mice. At six months, DICE and APP/PS1 mice exhibited similar plaque counts; however, DICE mice had significantly more plaques that were smaller in size. This could indicate either an increase in plaque formation that is not yet reflected by a higher total number at the chosen time-point in our study, or a change in the shape and density of the plaques. Plaque geometry is shaped by local proteostasis, neuritic dystrophy, and glial engagement (Xu et al., 2020). The smaller plaques in DICE mice, together with elevated GFAP and IBA1, suggest that epigenetically altered glia can remodel the peri-plaque environment, potentially enhancing compaction or modifying phagocytic processing without reducing plaque initiation (Tsering and Prokop, 2024). Moreover, this study is consistent with recent findings from Ramamrthy et al., 2023 which demonstrate that microglia exhibit acetylation changes linked to aging rather than to amyloid-beta plaque accumulation, whereas oligodendrocytes exhibit acetylation changes linked to amyloid-beta plaque load (Ramamurthy et al., 2022). Together, these findings underscore the important influence of the epigenome on glial function during aging and in age-related diseases, such as AD.

Our molecular findings complement the observed behavioral and histopathological evidence. GFAP expression was significantly higher in DICE relative to APP/PS1 mice, indicating that DNA damage exacerbates astrocytic activation in the context of amyloid plaques. These results are congruent with findings showing that astrocytes, upon sensing DNA damage or other cellular distress signals, can become activated, leading to a change in their morphology and function (Reyes-Ábalos et al., 2024; Wang et al., 2023). These findings potentially highlight astrocytes as a promising target for therapeutic intervention in AD. By better understanding the role of astrocytes in disease progression, especially in relation to DNA damage and cellular homeostasis, new strategies could be developed to slow or modify the course of AD (Kim et al., 2024).

The ICE system, which induces low level DSBs without high mutational burden, accelerates aging-associated epigenetic drift and supports the concept that chromatin remodeling and repair alone are sufficient to erode epigenetic information (Yang et al., 2023). Introducing this paradigm into an amyloid prone background demonstrates that recurrent, low level DNA damage can reshape AD trajectories at behavioral, inflammatory, and plaque morphological levels, emphasizing the significant impact that DNA damage has on the development and progression of AD.

Taking into consideration our findings, there are some alternative approaches that we would like to highlight for future studies. First, behavioral analyses, plaque morphology, and gliosis were assessed in ∼6 months-old DICE mice and given our findings employing longitudinal analyses could reveal whether early shifts in plaque geometry forecast later neurodegeneration or network dysfunction. Second, in our study DNA damage was induced for a 4-week period and given our findings, it would be interesting to explore models with prolonged or intermittent DNA damage to determine whether the timing and duration of genomic instability exert differential effects on AD pathology. Additionally, while the tamoxifen-I-*Ppo*I system produces dispersed DSBs with limited off-target mutagenesis, repair pathway choice and chromatin context might vary across cell types; single-cell multi-omic profiling (ATAC-seq plus CUT&Tag/CUT&RUN) in neurons, astrocytes, and microglia will be important to map the exact epigenetic drifts that accompany behavioral and histopathological changes. It would also be interesting to more deeply perform comparative analysis of sex-specific responses to the DSB-induced drift, given the observed sex differences in frailty and weight. Lastly, direct tests of causality between epigenetic states and glial function, for example, inducing DNA damage selectively in astrocytes/microglia on the DICE background, could determine whether heightened gliosis is a driver of the morphological differences or a response to them.

The DICE model reinforces the concept that DNA damage can accelerate the erosion of epigenetic information, consistent with the RCM hypothesis and the Information Theory of Aging. These findings suggest that AD progression may, in part, reflect noise in the epigenetic landscape that disrupts glial and neuronal identity. Future work should determine whether restoring youthful epigenetic information, via transient expression of OSK factors or chemical reprogramming, can reverse or mitigate amyloid-associated neuroinflammation.

In summary, in this study we applied controlled DNA damage in a genetically susceptible amyloid model and demonstrated that epigenetic erosion via DSBs modulates plaque morphology, amplifies gliosis, and exacerbates select behavioral phenotypes early in AD pathology. These findings thus support a model in which preservation of epigenomic integrity may represent a tractable strategy to modify AD progression.

## Materials and Methods

### Animals

Female and male hemizygous DICE mice and control groups (CTR, APP/PS1, ICE) were obtained by crossing hemizygous HA-ER^T2^-I-*Ppo*I/Cre-ER^T2^ female (“007001 - UBC-Cre-ERT2 Strain Details,” n.d.; Kim et al., 2016), and APP/PS1 male mice (“034832 - APP/PS1 Strain Details,” n.d.) (Jackson Laboratories, Bar Harbor, ME, USA). Animal genotypes were confirmed using polymerase chain reaction (PCR). Mice were given *ad libitum* access to food (Teklad Global Soy Protein-Free [Irradiated] type 2920X, Envigo, Indianapolis, IN, USA) and water, were group-housed by sex in ventilated caging (NexGen Mouse 500, Allentown Inc., Allentown, PA, USA) at the University of Rhode Island with no more than 5 mice/cage, and were kept on a 12:12 light:dark cycle at ∼22 °C and 30–70% humidity. Appropriate procedures were conducted to ensure animal safety and well-being. This study was conducted in line with the ethical standards and according to the Declaration of Helsinki and national and international guidelines and has been approved by the institutional review board at the University of Rhode Island.

### Tissue Preparation

Six-month-old CTR (WT or CRE), APP/PS1 (APP/PS1 or APP/PS1/CRE), ICE, and DICE mice were euthanized with 200 mg/kg sodium pentobarbital followed by cervical dislocation. Organs were collected and weighed accordingly, and rapidly frozen on dry ice and stored at -80°C until use or post-fixed in 10% formalin (Epredia, Portsmouth, NH, USA) at 4°C overnight, followed by submersion in 30% sucrose in 0.1M phosphate buffer. Alternatively, some mice were perfused with 10% formalin under anesthesia, and tissues were collected and post-fixed in 10% formalin at 4°C overnight, followed by submersion in 30% sucrose in 0.1M phosphate buffer. Tissues were embedded (Tissue-Plus OCT compound, Fisher Scientific, Waltham, MA, USA) and stored accordingly in -80°C until use.

### Cell Culture

Primary fibroblast cells were isolated from 6-month-old CTR, ICE, APP/PS1, and DICE mice. Ear biopsies were collected after euthanasia, cleaned with 70% ethanol, and rinsed three times in sterile PBS (SH30256.01, Cytiva, Marlborough, MA, USA). Excess hair was removed, and tissues were minced (1-3 mm) in sterile PBS and incubated in 0.25% trypsin-EDTA (25200056, Gibco, Waltham, MA, USA) at 37°C for 30 min. After digestion, minced tissues were maintained at 37°C and 5% CO_2_ in cell growth medium (DMEM with high glucose [4.5 g/L], containing 4 mM L-glutamine (1514022, Gibco, Waltham, MA, USA) and 1% penicillin/streptomycin (15140163, Gibco, Waltham, MA, USA) and supplemented with 20% FBS (89510-186, Avantor, Radnor Township, PA, USA) until cells have spread out from the tissue. Thereafter, cells were maintained in DMEM with FBS concentration lowered to 10%.

### Tamoxifen Treatment

All mice were fed a modified rodent diet (AIN-93G, Dyets Inc., Bethlehem, PA, USA) with 360 mg/kg Tamoxifen citrate (LSACT9262-1G, Wilkem Scientific, West Warwick, RI, USA) from 12 to 16 weeks of age and monitored for health and wellness daily. Tamoxifen citrate was delivered through food rather than intraperitoneal injection to ensure sustained nuclear HA-ER^T2^-I-*Ppo*I translocation and prolonged DNA damage (Yang et al., 2023).

Primary fibroblasts were treated with media containing 0.5 μM 4-Hydroxytamoxifen (4-OHT) (CAS 68047-06-3, Sigma Aldrich, St. Louis, MO, USA) for 0 and 72 h to induce DNA damage by activating the HA-ER^T2^-I-*Ppo*I endonuclease. Media containing 4-OHT was replenished every 24 hours to ensure adequate activation.

### Behavioral Experiments

All mice underwent a series of behavioral tests at 22 weeks of age, conducted in order of invasiveness: open-field, light/dark preference, elevated plus maze, Y-Maze, Barnes maze, social interaction, startle reflex, and pre-pulse inhibition. Behavioral tests were performed in a neutral and quiet environment and were conducted by the same researcher between 9:00 and 17:00 (light phase). Mice were acclimated for 1 h in a testing room at ∼22 °C, 30–70% humidity, and under low light (<100 lux). After each experimental trial, the apparatus was extensively cleaned with 70% ethanol to eliminate olfactory cues.

### Open-Field Assay

Spontaneous locomotor activity was measured using a square-shaped and darkened transparent chamber (40 × 40 × 30 cm) containing a base, four walls, and a lid with a grid of infrared beams at floor level and 7.5 cm above (Superflex Open Field, Omnitech Electronics, Columbus, OH, USA). The apparatus consisted of eight chambers allowing for eight mice to be tested at one time. Mice were placed in the center of the darkened chamber and left to explore the apparatus for 90 min while movements were monitored in 5-min intervals in the x-, y-, and z-planes. Distance traveled, duration, rearing, bouts, rest time, and velocity were recorded and analyzed (Fusion v6.5 software system, Omnitech Electronics, Columbus, OH, USA) in peripheral and center locations of the arena.

### Light/Dark Preference Test

Using the Superflex Open Field system (Omnitech Electronics, Columbus, OH, USA), animals were placed into a chamber that was divided into two sections of equal size by a partition with an opening such that one half of the chamber was brightly illuminated and the other half of the chamber was dark. Mice were placed in the center of the brightly illuminated side of the apparatus and left to explore the apparatus for 30 min. Distance traveled, duration, rearing, bouts, rest time, vertical movement, and velocity were recorded and analyzed (Fusion v6.5 software system, Omnitech Electronics, Columbus, OH, USA) in both light and dark compartments.

### Elevated Plus Maze

Anxiety and cognition were assessed using an elevated-plus maze apparatus (height 59 cm x width 7 cm x length 30 cm with a wall height of 15 cm) containing a cross-shaped platform with four arms (two open arms and two enclosed arms) as previously described ((Gaspar et al., 2023); CleversSys Inc., Reston, VA, USA). Briefly, each mouse was transported to and from the maze in a darkened container to minimize external stress and was placed in the center of the maze and allowed 5 min to explore the apparatus at ∼300 lux. Measures such as duration, distance traveled, and number of bouts in the closed and open arms were recorded and evaluated via a camera tracking system (ANY-maze, Stoelting, Wood Dale, IL, USA).

### Y-Maze

Short-term memory was assessed using a Y-maze (30 x 7 x 15 cm; CleversSys Inc., Reston, VA, USA), consisting of three symmetrical arms, one of which could be closed. Mice were transported to and from the maze in a darkened container to minimize external stress and stimuli. In the first trial, mice explored two open arms for 3 min at ∼300 lux, with the third arm closed. After a 15-min rest in the home cage, mice underwent a second 3-min trial with the previously closed arm (novel arm) opened. Time spent, distance traveled, and number of entries into the novel arm were recorded via a camera tracking system (ANY-maze, Stoelting, Wood Dale, IL, USA).

### Barnes Maze

Spatial learning and memory were evaluated using an apparatus made from a circular, gray, non-reflective, powder-coated metal base platform with a diameter of 91 cm (Stoelting Co., Kiel, Wisconsin, USA). The apparatus contained 20 holes lining the periphery of the apparatus, each with a diameter of 5 cm, surrounded the perimeter of the maze at 2.5 cm from the edge. Of the 20 holes, one hole was designated as the escape location and a goal box (height 7.5 cm x width 5 cm x length 15 cm) was situated directly beneath the hole to allow the animal to escape from the open apparatus. Four visual cues (large black circle, square, triangle, and a cross) were placed evenly surrounding the maze and a bright light (∼800 lux) above the maze was used as a deterrent to encourage the mouse to escape from the open apparatus. The test was performed over the course of five days and consisted of four consecutive training days, with four trials performed each day, followed by a probe test on the fifth day. During training days, the mouse was placed under a plastic, opaque acclimation container in the center of the maze for 15 s. Once the container was removed, the animal was given 3 min to locate the escape box. If it did not find the escape box within time allowed, the animal was gently guided to and left inside the escape box for 1 min. After each trial, mice were returned to their home cage for 15 min. During the probe day, the mouse was again placed under a plastic, opaque acclimation container in the center of the maze for 15 s and given 90 s to explore the apparatus and locate the escape box that is now removed. Parameters such as duration spent near the escape box, bouts in the escape hole, and duration spent in each quadrant were monitored via a camera tracking system (ANY-maze, Stoelting, Wood Dale, IL, USA).

### Social Interaction

Sociability was measured using the Superflex Open Field system fitted with two grid enclosures in each chamber (Omnitech Electronics, Columbus, OH, USA). Using this test, mice underwent two phases: habituation and testing. During the first phase (habituation), each mouse was placed in the center of the chamber and left to habituate to the apparatus for 10 min until returned to their home cage. Immediately after habituation, the same mouse underwent a second phase (testing) and was placed in the arena with two grid enclosures, one containing a Lego block and the other one containing a sex-matched “stranger” mouse, and was allowed to explore for 10 min. Parameters such as duration spent and number of bouts near each enclosure was measured and analyzed (Fusion v6.5 software system, Omnitech Electronics, Columbus, OH, USA).

### Startle Reflex

The acoustic startle reflex was measured using the startle reflex behavioral test. The apparatus consisted of an isolation cabinet (height 46 cm x width 38 cm x length 36 cm) containing a plexiglass tube (length 9 cm with 3 cm diameter) glued to a plastic plate containing an accelerometer underneath (SR-LAB-Startle Response System, SD Instruments, San Diego, CA, USA). Mice were placed inside of the plexiglass tube and allowed 5 min to acclimate to the apparatus. During the test, mice underwent three trials of randomly spaced acoustic noises (20– 120 dB), with each mouse exposed randomly to each acoustic level a total of six times. Parameters such as the maximum millivolt of startle were measured and analyzed in relation to acoustic sound intensity.

### Pre-Pulse Inhibition

The ability to attenuate a startle reflex response was measured using the pre-pulse inhibition behavioral assay using the startle reflex apparatus previously described (SR-LAB-Startle Response System, SD Instruments, San Diego, CA, USA). During the test, mice underwent exposures to the startle alone, then 10 blocks with 6 different types of trials. These trials included a null trial where no stimuli were presented, startle trial of 120 dB, startle plus pre-plus with variable dB and pre-pulse alone. The trials were randomly interspersed, and each trial started with a 50 ms null period to monitor movements at baseline followed by exposure to the pre-pulse alone and that response was recorded. Following this, the startle stimuli were presented, and the responses recorded. Parameters such as the maximum millivolt of startle with each pre-pulsed stimulus were measured and analyzed in relation to acoustic sound intensity.

### Frailty Index Assessment

Aging phenotype was scored using the frailty index (FI) assessment developed previously by Whitehead et al., 2014 (Whitehead et al., 2014). FI was assessed in all mice by the same researcher, blinded to genotype, after completion of all behavioral tests. Body weight was recorded, and body temperature was measured three times near the anus using an infrared thermometer, then averaged. Scores of 0, 0.5, or 1 were assigned based on deviation from the group average for weight and temperature. Additionally, 29 items related to integument, musculoskeletal, vestibulocochlear/auditory, ocular/nasal, digestive/urogenital, respiratory systems, and overall discomfort were scored on the same scale (0 = least severe, 1 = most severe). FI was calculated by dividing the total score by the number of items assessed.

### Histology

Tissues were cryosectioned at -21°C (Leica BioSystems, CM1950, Wetzlar, Germany) at 30 μm and either mounted directly onto slides (VWR Colorfrost Plus, Radnor, PA, USA) or collected in netwells (3477, Corning, Corning, NY, USA) containing PBS for processing using the free floating method (Potts et al., 2020). For Thioflavin S and cresyl violet staining, free-floating sections were mounted onto slides accordingly, hydrated through a series of decreasing concentrations of ethanol (100%, 95%, and 75%), stained with 1% Thioflavin S solution (T1892, Sigma Aldrich, St. Louis, MO, USA) for 1 h or 0.1% Cresyl violet acetate solution (26089-01, Electron Microscopy Sciences, Hatfield, PA, USA) for 3-4 min at RT, further dehydrated in increasing concentrations of ethanol (70%, 95%, and 100%), and mounted with Entellan (1.07960, Sigma Aldrich, St. Louis, MO, USA). For 1-42 A*β,* GFAP, and IBA1 visualization, floating sections were washed three times with TBS and blocked in 3% horse serum in TBS + 0.3% Triton-X-100 solution. Sections were washed three times with TBS and incubated in primary antibody (1-42 A*β*: 1:500, H31L21, Invitrogen, Waltham, MA, USA; GFAP: 1:1000, PA514358, Thermo Scientific, Waltham, MA, USA; IBA1: 1:500, 011-27991, FujiFilm Irvine Scientific Inc., Santa Ana, CA, USA) overnight at 4°C, washed three times with TBS, and incubated in secondary antibody at RT (Alexa 488: 1:500, 711-545-152, Jackson ImmunoResearch Laboratories Inc., West Grove, PA, USA; Alexa 594: 1:500, 705-585-003, Jackson ImmunoResearch Laboratories Inc., West Grove, PA, USA). After washing two times with TBS, nuclei were stained with Hoechst (1:2000, H1399, Invitrogen, Waltham, MA, USA) incubated in PBS (pH = 7.4) for 15 min, mounted accordingly onto slides, and coverslipped (CS-24X50, AmScope, Irvine, CA, USA) using antifade aqueous mounting medium.

### Microscopy and Amyloid-Beta Quantification

Bright-field and fluorescence imaging were conducted to visualize and record immunofluorescence and staining results (Leica THUNDER DMi8 3D inverted epifluorescence microscope and LAS X 3D Analysis Software v. 2018.7.3, Leica Microsystems, Wetzlar, Germany). Images were processed using appropriate software ((Schindelin et al., 2012); FIJI v2.1.0/1.53c, Madison, WI, USA). Fluorescent amyloid-beta plaques were quantified manually by the same blinded researcher, and the number of amyloid-beta plaques were counted two times, averaged, and normalized to the number of hippocampal sections for each mouse. Amyloid-beta plaque area and perimeter were determined using appropriate software ((Schindelin et al., 2012); FIJI v2.1.0/1.53c, Madison, WI, USA) as previously reported (Christensen and Pike, 2020).

### Western Blot Analysis

To detect HA-ER^T2^-I-*Ppo*I protein expression to validate the ICE system, cells were rinsed three times with cold PBS and lysed in RIPA Buffer (50 mM Tris-HCl pH 7.4, 150 mM NaCl, 0.5% deoxycholic acid, 0.1% sodium dodecyl sulfate, 2 mM EDTA, 1% Triton X-100) containing protease inhibitors (78438, Thermo Scientific, Waltham, MA, USA). To detect expression of hu-APP and GFAP in brain, cerebella were lysed in RIPA buffer containing protease inhibitors. Cell or tissue lysates were briefly sonicated (QSonica, Newtown, CT, USA) and centrifuged at 10^4^ × g for 10 min. The supernatant was then transferred to a clean Eppendorf tube, and protein concentrations were quantified using a BCA-based assay kit (23225, Thermo Scientific, Waltham, MA, USA). Equal amounts of protein (10–50 μg) were separated by electrophoresis using as SDS-PAGE gel (4-20% or 8-16%; BioRad, Hercules, CA, USA) and blotted onto 0.2 or 0.45 μm nitrocellulose membranes (10600002, Cytiva, Marlborough, MA, USA). Membranes were blocked using 5% milk in 0.1% TBS-Tween solution and incubated with primary antibodies (HA-Tag: 1:1000, C29F4, Cell Signaling, Danvers, MA, USA; hu-APP: 1:1000, 803004 BioLegend, San Diego, CA, USA; *β*-Actin: 1:50,000, A3854, Sigma Aldrich, St. Louis, MO, USA) in 5% milk in 0.1% TBS-Tween solution overnight at 4°C. After washing three times in 0.1% TBS-Tween, membranes were incubated for 1 h at RT with secondary antibody HRP-conjugated (1:3000, 1706515, Bio-Rad) in 5% milk in 0.1% TBS-Tween solution. Immunocomplexes were visualized using ECL solution (Clarity ECL, Bio-Rad, Hercules, CA, USA) via chemiluminescence (ChemiDoc, Bio-Rad, Hercules, CA, USA). All samples were first normalized to β-actin and subsequently expressed relative to the APP/PS1 group.

### Sanger Sequencing

Clean DNA was extracted from olfactory bulbs of 6-month-old CTR, APP/PS1, and DICE mice with and without tamoxifen treatment. Briefly, tissue was incubated in buffer containing 10 mM Tris-HCl (pH = 8.0), 25 mM EDTA, 10% SDS, 100 mM NaCl, and 200 µg/mL proteinase K solution overnight. The next day, samples were centrifuged, and supernatant was mixed with equal volume of 25:4:1 phenol/chloroform/isoamyl alcohol (25:4:1 PCIAA, P2069, Sigma Aldrich, St. Louis, MO, USA). Samples were centrifuged, and the aqueous phase was collected and subjected to an additional chloroform-only extraction to further remove residual phenol and protein contaminants. DNA was then precipitated using 3 M sodium acetate and isopropyl 100% alcohol. The sample was maintained at −20°C for 20 min to facilitate DNA precipitation. DNA precipitate was washed one-time with 70% ethanol and resuspended with tris-EDTA buffer. DNA concentration and purity were confirmed (NanoDrop, ND-2000, Thermo Scientific, Waltham, MA, USA), and hu-APP CDS sequence amplified with primers (Thermo Scientific, Waltham, MA, USA) specific to hu-APP (hu-APP_For_: GAACCATTTCAACCGAGCTGAAGC; hu-APP_Rev_: TTAGTTCTGCATTTGCTCAAAGAA) using a high-fidelity DNA polymerase (Phusion^™^, New England Biolabs, Ipswich, MA, USA). The amplified fragments were separated on a 1% agarose gel and purified (E.Z.N.A. Cycle Pure Kit, Omega Bio-Tek, Norcross, GA, USA). Samples were then sequenced (3500xL Genetic Analyzer, Applied Biosystems, Foster City, CA, USA). The primer sequences (Thermo Scientific, Waltham, MA, USA) utilized were as follows: hu-APP_For1_: GAACCATTTCAACCGAGCTGAAGC; hu-APP _For2_: ATGTTTGTGAGACCCATCTTCACT; hu-APP_For3_: AAGTACCTGGAGACACCCGGGGAC; hu-APP_For4_: GATCTACGAGCGCATGAACCAGTC; hu-APP_For5_: GTTCAAACAAAGGCGCCATCATCG; hu-APP_Rev1_: TTAGTTCTGCATTTGCTCAAAGAA; hu-APP_Rev2_: CCCCATTCACGGGAAGGAGCTCCA; hu-APP_Rev3_: TGCTGTCTCTCATTGGCTGCTTCC; hu-APP_Rev4_: GTAGTCTGTGTCCGCTCCACCCCA; hu-APP_Rev5_: CTGACGGGTCTGACTCCCACTTTC.

### HA-I-PpoI Immunoprecipitation

5-alpha competent E. *coli* bacteria (C2988, New England Biolabs, Ipswich, MA, USA) heat-shock transformed with pICE-HA-NLS-I-*Ppo*I plasmid (#46963, Addgene, Watertown, MA, USA; (Britton et al., 2013)) were grown overnight in liquid Luria Broth (LB), and plasmid isolated and purified by miniprep (E.Z.N.A. Plasmid DNA Mini Kit I Omega Bio-Tek, Norcross, GA, USA). DNA concentration and purity wereassessed using a spectrophotometer (NanoDrop, ND-2000, Thermo Scientific, Waltham, MA, USA). Human U-2 OS cells were grown at 37°C and 5% CO_2_ in cell growth medium (DMEM with high glucose [4.5 g/L], containing 4 mM L-glutamine (1514022, Gibco, Waltham, MA, USA) and 1% penicillin/streptomycin (15140163, Gibco, Waltham, MA, USA)) supplemented with 10% FBS (89510-186, Avantor, Radnor Township, PA, USA), and subsequently transfected with pICE-HA-NLS-I-*Ppo*I plasmid. After two days cells were rinsed three times with cold PBS and lysed in non-denaturing lysis buffer (20 mM Tris-HCl, 136 mM NaCl, 2 mM EDTA, 0.01% Triton-X-100, and 12% Glycerol) containing protease inhibitors (78438, Thermo Scientific, Waltham, MA, USA). The cell lysate was briefly sonicated (QSonica, Newtown, CT, USA) and incubated with Pierce Anti-HA agarose beads (26182, Thermo Scientific, Waltham, MA, USA) overnight at 4°C. The next day, beads were collected by centrifugation and washed three times with TBS solution containing 0.05% Tween-20, and finally resuspended in I-*Ppo*I digestion buffer (6mM Tris-HCl, 100 mM NaCl, 1 mM DTT, 6 mM MgCl_2_, solution pH = 7.5; (McCarthy and Romanowski, 2006)). Active immunoprecipitated HA-NLS-I-*Ppo*I was maintained at 4°C before assay. To confirm HA-I-PpoI isolation, beads were boiled in 4X Laemmli sample loading buffer (1610747, BioRad, Hercules, CA, USA), separated on SDS-PAGE gel (4-20%, BioRad, Hercules, CA, USA), and blotted onto 0.45 μm nitrocellulose membrane (10600002, Cytiva, Marlborough, MA, USA). The membrane was blocked using 5% milk in 0.1% TBS-Tween solution and incubated with HA-Tag primary antibody (1:1000, C29F4, Cell Signaling, Danvers, MA, USA) in 5% milk and 0.1% TBS-Tween solution overnight at 4°C. After washing three times in 0.1% TBS-Tween, the membrane was incubated for 1 h at RT with secondary antibody HRP-conjugated (1:3000, 1706515, Bio-Rad, Hercules, CA, USA) in 5% milk and 0.1% TBS-Tween solution. HA-NLS-I-*Ppo*I immunocomplex was visualized (Clarity ECL, Bio-Rad, Hercules, CA, USA) via chemiluminescence (ChemiDoc, Bio-Rad, Hercules, CA, USA).

### HA-I-PpoI Digestion

A series of restriction enzyme digestion reactions were completed to confirm that there was no recognition site on the hu-APP gene for the I-*Ppo*I endonuclease. DNA concentration was amplified with primers (Thermo Scientific, Waltham, MA, USA) specific to hu-APP (hu-APP_For_: GAACCATTTCAACCGAGCTGAAGC; hu-APP_Rev_: TTAGTTCTGCATTTGCTCAAAGAA) using PCR with a high-fidelity DNA polymerase (Phusion^™^, New England Biolabs, Ipswich, MA, USA), and DNA fragment size, concentration and purity confirmed (NanoDrop, ND-2000, Thermo Scientific, Waltham, MA, USA). A positive control plasmid p201B/Cas9 (#59177, Addgene, Watertown, MA, USA (Jacobs et al., 2015)) was used to confirm the proper activation of immunoprecipitated I-*Ppo*I endonuclease as described above. The p201B/Cas9 plasmid along with the hu-APP PCR product underwent a series of three digestion reactions overnight in a 37°C incubator (Genie Temp-Shaker 300, LabRepCo, Horsham, PA, USA): a reaction with no restriction enzyme, a reaction with immunoprecipitated I-*Ppo*I endonuclease, and a reaction with immunoprecipitated I-*Ppo*I endonuclease with an additional restriction enzyme confirmed to have a recognition site for that plasmid sequence (#59177, Addgene, Watertown, MA, USA (Jacobs et al., 2015); AgeI-HF [p201B/Cas9] and BbSI-HF [hu-APP], New England Biolabs, Ipswich, MA, USA). All digestion reactions were mixed with DNA loading buffer and separated on a 1% agarose gel for visualization (ChemiDoc, Bio-Rad, Hercules, CA, USA).

### RNA Extraction and Quantitative PCR (qPCR)

RNA was extracted from olfactory bulbs from 6-month-old CTR, APP/PS1, and DICE mice (R2073-A, Direct-zol RNA MiniPrep Plus with TRI Reagent kit, Zymo Research, Irvine, CA, USA). RNA concentration and purity were assessed (NanoDrop, ND-2000, Thermo Scientific, Waltham, MA, USA) and all samples were normalized to the concentration of the most dilute sample in the set. cDNA synthesis was performed (E3010, Lunascript RT Supermix Kit New England Biolabs, Ipswich, MA, USA). Successful reverse transcription was confirmed by standard PCR using primers specific for β-actin. Reactions were carried out (M3003, Luna qPCR SYBR Green Mastermix, New England Biolabs, Ipswich, MA, USA) using an qPCR instrument (Roche LightCycler 480, Roche Holding AG, Basel, Switzerland). Data were analyzed (LightCycler software, Roche Holding AG, Basel, Switzerland). The primer sequences (Thermo Scientific, Waltham, MA, USA) used were: mut-APP_For_: 5′-AAGAGATCTCTGAAGTGAATC-3′, mut-APP_Rev_: 5′-TTGTTTGAACCCACATCTTCT-3′; *β*-Actin_For_: 5′-ACCTTCTACAATGAGCTGCG-3’; *β*-Actin_Rev_: 5’-CTGGATGGCTACGTACATGG-3’.

### Statistics

Data are presented as mean value (M) with SEM and sample size (N) indicated in the figure legends. Statistical analyses were performed using appropriate software (Microsoft Excel, Redmont, WA, USA and GraphPad Prism v. 9, San Diego, CA, USA) by applying either a two-way ANOVA using Tukey multiple comparison or unpaired t-test, with an α level of 0.05.

Significances are denoted in figures with \**p* < 0.05, ** *p* < 0.01, *** *p* < 0.001, **** *p* < 0.0001, and trending as an α *p* < 0.10. Normality was tested by using the D’Agostino & Pearson test and the Shapiro-Wilk test. Outliers were determined using the outlier calculator with an α level of 0.05 (GraphPad Prism v. 9, San Diego, CA, USA).

## Acknowledgements

We thank Dr. David Sinclair and Dr. Philipp Oberdoerffer for the use of the ICE mouse model. We are grateful for the assistance of the animal care staff at the University of Rhode Island. Figures 1A, B, D and 2B were created using BioRender.

## Author Contributions

This publication encompasses the thesis of Sydney Bartman. J.M.R. and G.C. conceived the project and J.M.R. provided funding. D.A.S. developed the ICE mouse model and provided intellectual input on its use for this study, helped design experiments, and edited the manuscript.

G.C. and J.M.R. provided supervision. S.B., G.C. and J.M.R. designed experiments, S.B. performed all experiments and analyzed the data, and D.H., D.Z., L.G., and H.T-W. assisted in performing some experiments. S.B., G.C. and J.M.R. wrote the manuscript and all authors have read and agreed to the published version of the manuscript.

## Institutional Review Board Statement

The animal study protocol (#AN1920-020) was approved by the Institutional Review Board of University of Rhode Island under G.C. and J.M.R.

## Informed Consent Statement

Not applicable.

## Data Availability Statement

Data generated from this study are available upon request.

