## Supplementary Material for "DNA Break-Induced Epigenetic Alterations Promote Plaque Formation and Behavioral Deficits in an Alzheimer’s Disease Mouse Model"

### Supplementary Figures

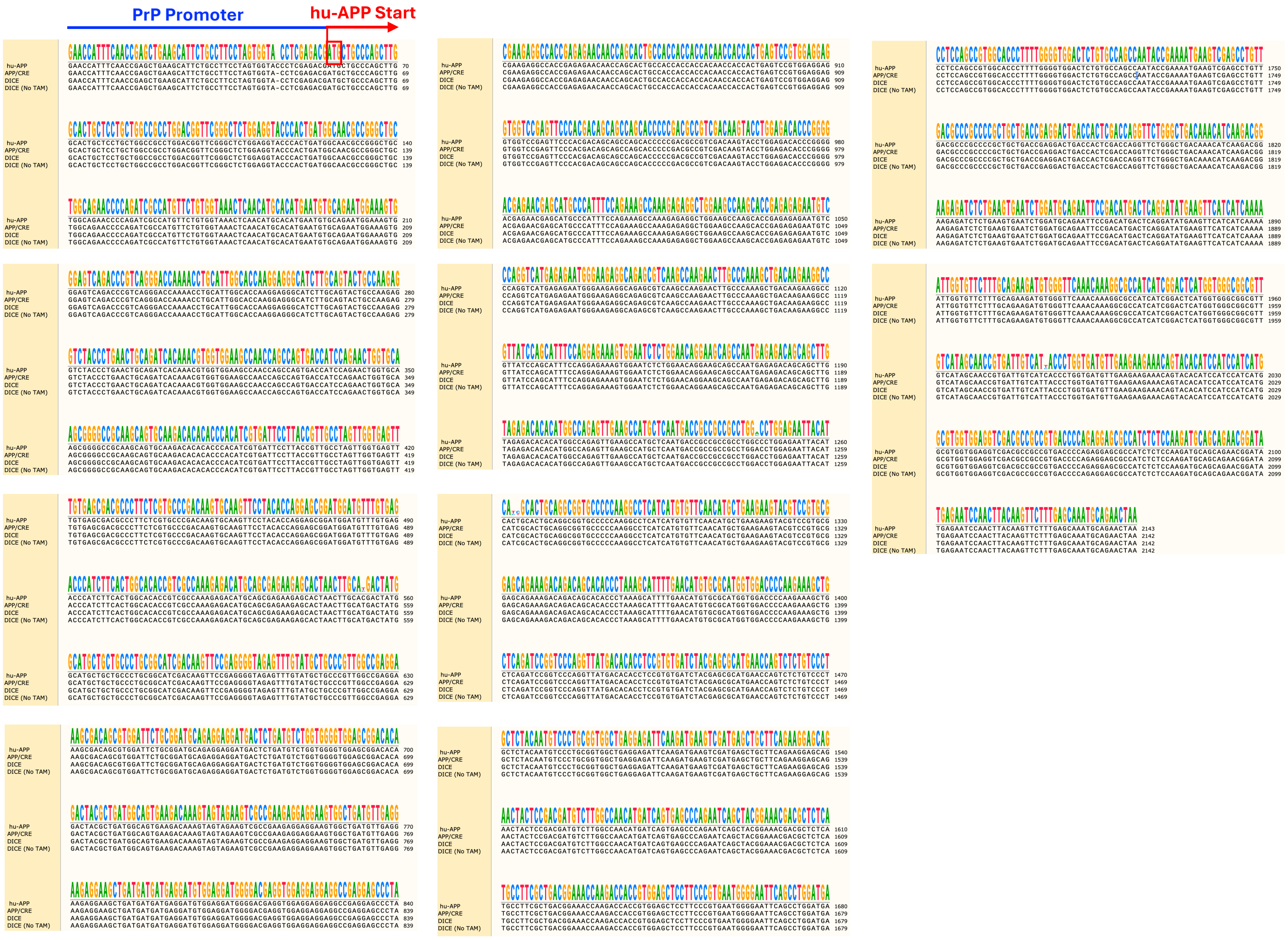


**Supplementary Figure 1. Sanger sequencing of the hu-APP gene in APP/PS1 and ICE mice.** Alignment of hu-APP gene sequences from APP/PS1, DICE mice treated with tamoxifen, and DICE mice not treated. Sequencing demonstrates the absence of mutations in the coding region.

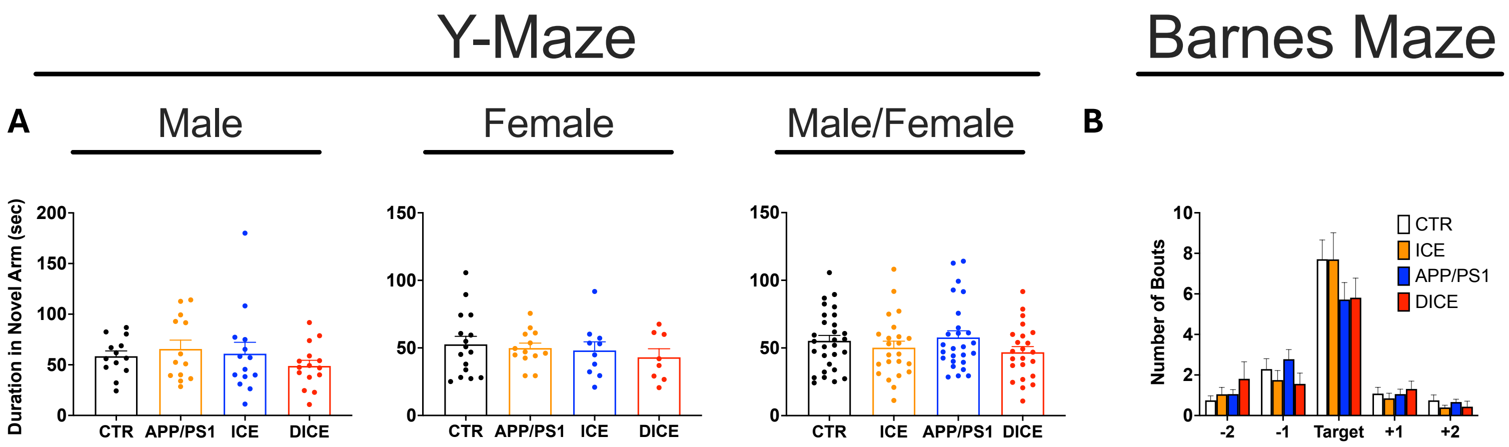


**Supplementary Figure 2. Y-maze and Barnes maze analyses indicate preserved working memory and spatial learning in DICE mice following tamoxifen treatment. (A)** Number of entries into the novel arm in the Y-maze. DICE mice made fewer entries than ICE mice but did not differ from APP/PS1 or CTR mice [CTR ($N_{Male}$=13; $N_{Female}$=16), ICE ($N_{Male}$=14; $N_{Female}$=10), APP/PS1 ($N_{Male}$=13; $N_{Female}$=13), and DICE mice ($N_{Male}$=15; $N_{Female}$=8)]. **(B)** Number of entries into the target hole in the Barnes maze. No significant differences were observed between male and female DICE mice and APP/PS1, ICE, or CTR controls [CTR ($N_{Male}$=9; $N_{Female}$=15), ICE ($N_{Male}$=10; $N_{Female}$=10), APP/PS1 ($N_{Male}$=10; $N_{Female}$=8), and DICE ($N_{Male}$=11; $N_{Female}$=5)]. Data are presented as mean ± SEM. Statistical analyses were performed using an unpaired t-test with Welch's correction; *a* indicates a trend p<0.10.

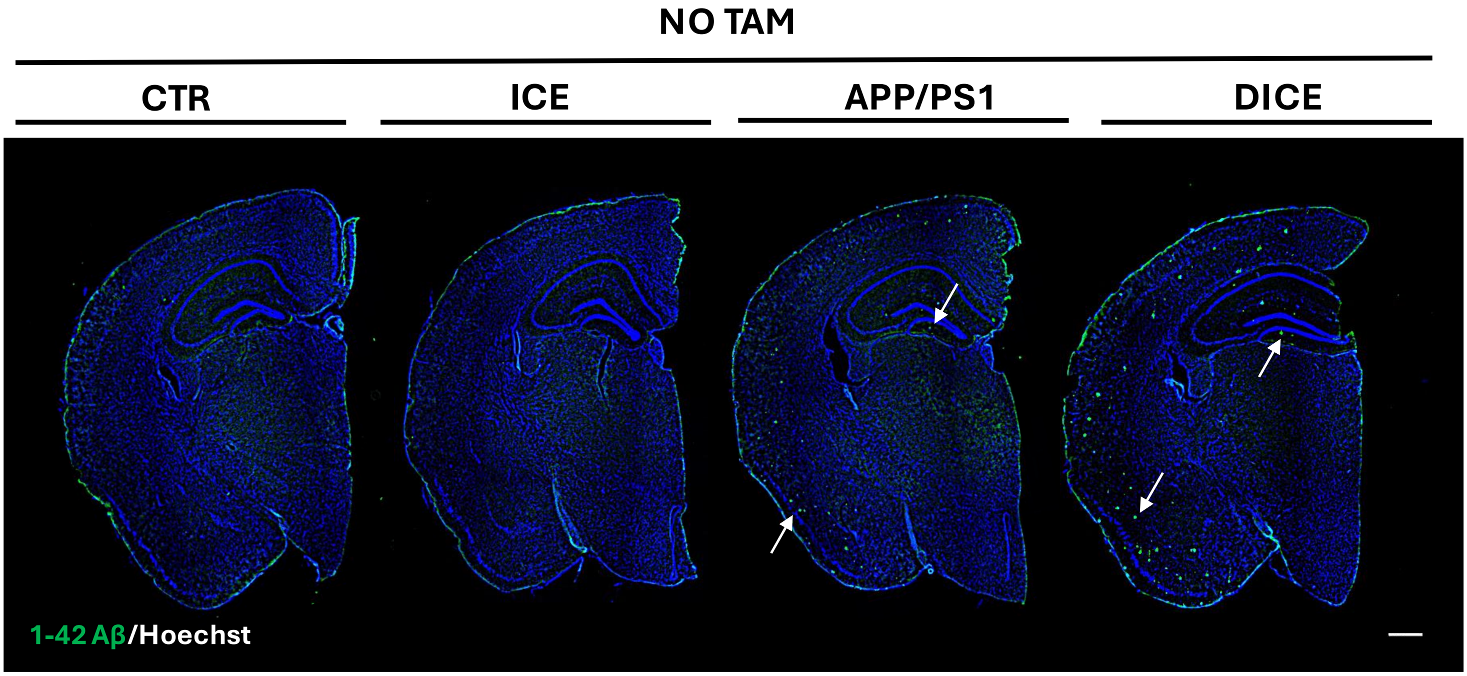


**Supplemental Figure 3. Tamoxifen treatment has no effect on amyloid-beta plaque formation in DICE and APP/PS1 mice.** Representative fluorescent immunohistochemistry for Aβ1-42 and Hoechst in coronal brain sections from 6-month-old CTR, ICE, APP/PS1, and DICE male and female mice (N=4 mice per group) that did not receive tamoxifen (5X; scale bar = 500 µm) shows no change in plaque formation in the absence of tamoxifen.

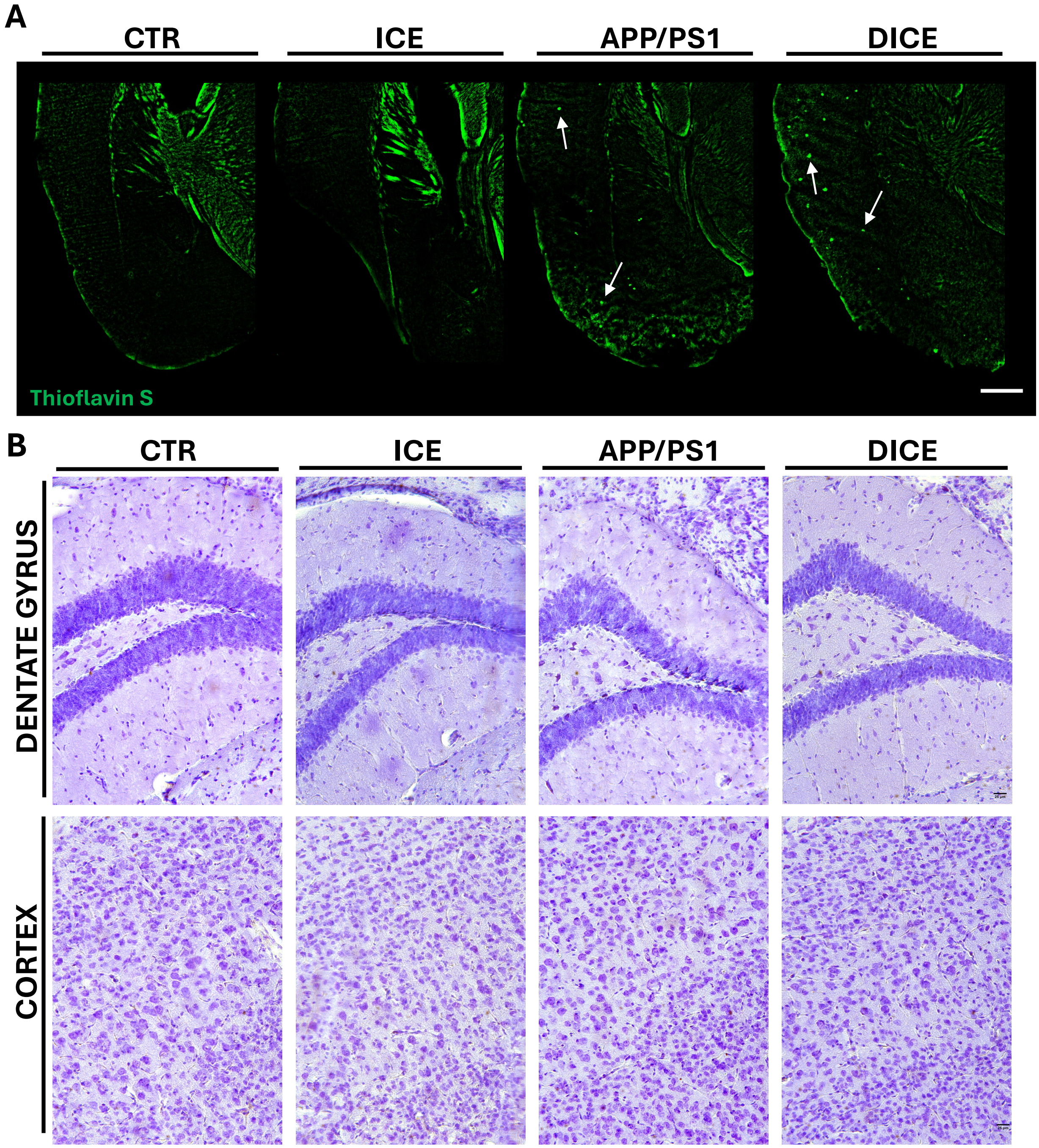


**Supplementary Figure 4. DICE mice show no significant changes in amyloid beta plaque load and Nissl body presence compared to APP/PS1 mice. (A)** Representative fluorescent immunohistochemistry for Thioflavin S in coronal brain sections from 6-month-old male and female CTR, ICE, APP/PS1, and DICE mice (N=3 mice per group) that did not receive tamoxifen treatment and show no changes in plaque formation in the absence of tamoxifen (5X; scale bar = 500 µm). Examples of plaques are indicated by white arrows. **(B)** Cresyl violet staining demonstrates no gross changes in morphology or detectable pyknotic bodies in dentate gyrus or cortex in CTR, ICE, APP/PS1, and DICE mice (N=2 per group) (20X; scale bar = 25 µm).
